# Numerosity adaptation reflects multiple levels of sensory processing

**DOI:** 10.64898/2026.07.21.739325

**Authors:** Tomoki J. Takahashi, Neil W. Roach, Masamichi J. Hayashi

## Abstract

Psychophysical and neuroimaging studies have repeatedly suggested that perceived numerosity is susceptible to adaptation, involving high-level number representations. However, whether adaptation to low-level visual features contributes to numerosity adaptation remains unclear. Here, we provide evidence that numerosity adaptation involves not only high- but also low-level visual processing stages. Through a series of psychophysical experiments, we found that the effect of numerosity adaptation was significantly reduced, but not eliminated, when the contrast polarity of stimuli was inverted relative to the preceding adaptor. Follow-up experiments confirmed that the persistence of the aftereffects was not due to retinal adaptation or top-down decision bias. Finally, a computational model incorporating both high- and low-level adaptation successfully reproduced the behavioral pattern. These findings suggest that numerosity adaptation involves at least two stages of visual processing: adaptation to low-level visual features at early sensory stages and to numerosity at higher levels of the visual processing hierarchy.

**Highlights:**

- Number adaptation is sensitive to mismatches in contrast polarity
- Mismatched stimuli reduce aftereffects but do not eliminate them
- Early sensory adaptation may modulate higher stage numerosity processing

## Introduction

Many animals, including humans, can estimate the number of objects in a scene without explicitly counting each item individually^1–7^. This ability plays a crucial role in decision-making across various contexts, such as assessing the number of allies or enemies in a group or evaluating the abundance of available food. In humans, several studies have shown that children’s ability to discriminate non-symbolic numbers reliably predicts both their current and future mathematical performance^8–12^, highlighting numerosity perception as a foundational skill for mathematical competence.

Past psychophysical studies have shown that perceived numerosity exhibits adaptation effects comparable to those reported for many other visual attributes, such as orientation, motion, size, and depth^13–17^. Specifically, a medium numerosity is perceived as less numerous following adaptation to a large numerosity, whereas it is perceived as more numerous following adaptation to a small numerosity (i.e., a negative aftereffect). Importantly, perceived numerosity is widely considered to be relatively robust to changes in low-level visual features, such as density, total area, and contrast, which often covary with numerosity^18–21^ and exert relatively minor influence on numerosity judgments^22–26^. Neuroimaging studies have implicated numerosity-tuned neural populations in high-level association cortices, including the parietal cortex, by demonstrating systematic biases in the population responses following adaptation to specific numerosities^27–29^. Together, these findings have reinforced the prevailing view that numerosity adaptation primarily reflects changes in high-level numerosity representations rather than adaptation to low-level image properties.

However, a recent psychophysical study has challenged this view. Grasso and colleagues^30^ reported that numerosity adaptation produced “virtually no [after]effect” when the color of the test stimulus differed from that of the preceding adaptor stimulus. This finding suggests that low-level visual features may play a more substantial role in numerosity adaptation than previously assumed. Such sensitivity to low-level features is potentially problematic because many conventional studies have used dot arrays containing equal numbers of black and white dots to control overall luminance, implicitly assuming that numerosity perception is insensitive to low-level visual features. Under this assumption, observers are expected to encode the total number of dots (e.g., six dots), rather than separately encoding subsets defined by contrast polarity (e.g., three black dots and three white dots).

The dependence of numerosity adaptation on low-level visual features can be explained by at least two possible mechanisms. One possibility is that high-level numerosity representations, such as numerosity-tuned neurons in association cortices, are themselves sensitive to low-level visual features (i.e., a direct effect). Alternatively, such sensitivity may arise indirectly from adaptation at early sensory-processing stages, such as the visual cortex, whereby adaptation to low-level visual features modulates the strength of input signals to otherwise feature-invariant numerosity representations (i.e., an indirect effect). These two hypotheses generate distinct predictions when the congruency of low-level visual features, specifically contrast polarity, between adaptor and test stimuli is manipulated in a numerosity adaptation paradigm. If the sensitivity is driven solely by a direct effect, numerosity aftereffects should be observed only when the low-level visual features of the adaptor and test stimuli are matched, and should be absent when they are incongruent. In contrast, if the sensitivity arises from an indirect effect, the numerosity aftereffect should be reduced but still present under incongruent conditions, because adaptation at early sensory stages would reduce, but not eliminate, the input reaching otherwise feature-invariant numerosity representations.

To distinguish between these hypotheses, we conducted a series of psychophysical numerosity adaptation experiments using dot arrays composed of positive, white, and negative, black, contrast polarities presented against a gray background. In the first experiment, we demonstrated that adaptation to dot arrays with mixed contrast polarities induces systematic biases in perceived numerosity relative to dot arrays with a single contrast polarity. We then systematically manipulated the match or mismatch in contrast polarity between adaptor and test stimuli and showed, across a series of experiments, that although the magnitude of the numerosity aftereffect was attenuated, robust aftereffects persisted even when contrast polarity was incongruent. Finally, we evaluated whether a computational model incorporating both early sensory adaptation and adaptation within numerosity representations could account for the observed behavioral pattern.

## Results

### Experiment 1: Low-level sensory features affect numerosity adaptation

To examine the influence of low-level sensory features on numerosity adaptation, our first experiment investigated whether numerosity adaptation with mixed contrast polarities of dot arrays, typically employed in previous numerosity adaptation studies, differs from adaptation with a single contrast polarity (i.e., all black or all white dots). The adaptor and probe stimuli were presented sequentially, each consisting of two dot arrays displayed in the left and right visual fields (Figure 1A). Participants were instructed to indicate which of the two dot arrays in the probe stimulus contained a larger number of dots.

**Figure 1.**
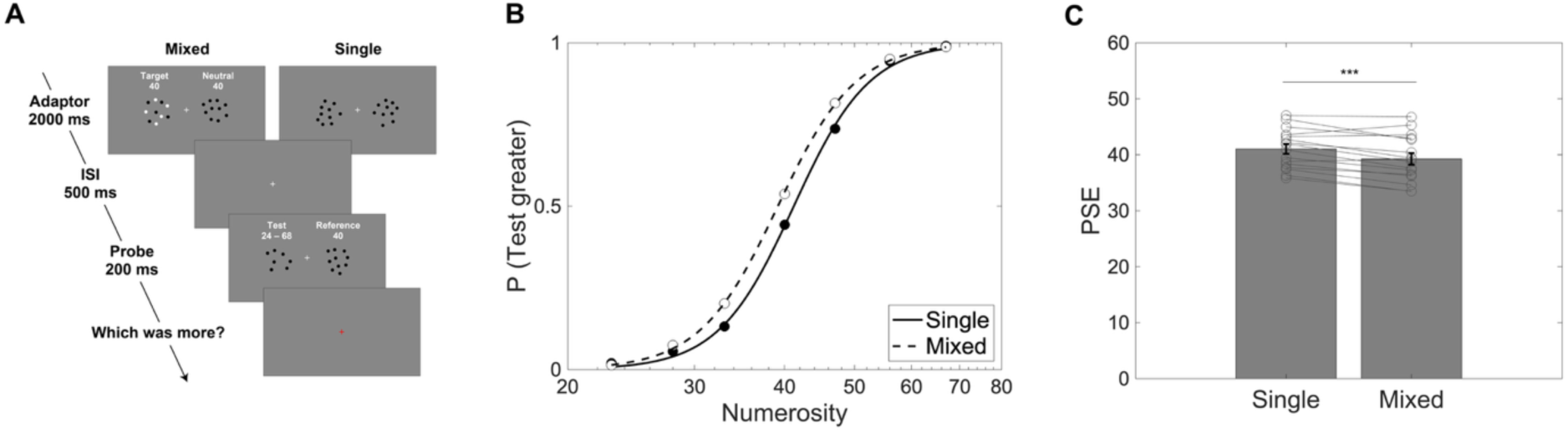
Stimulus sequence and results of Experiment 1. (A) In each trial, adaptor and probe stimuli were presented sequentially, each consisting of two dot arrays displayed in the left and right visual fields. During the response period (red fixation), participants responded which of the two dot arrays in the probe stimulus contained more dots. In the mixed-polarity condition, the target adaptor consisted of an equal number of black and white dots, whereas in the single-polarity condition all dots shared the same contrast polarity (either black or white), which matched that of the probe stimulus. For illustration purposes, fewer dots are shown than were used in the actual experiment. (B) Psychometric function fitted to the mean proportion of ‘test greater’ responses across participants for single-polarity (solid line) and mixed-polarity (dashed line) conditions. (C) Mean PSEs for the single-polarity and mixed-polarity conditions across participants. Gray circles represent individual participants’ data. Error bars denote standard error. *** p < 0.001.

The numerosity of the test stimulus in the probe varied on a trial-by-trial basis between 23 and 67 dots. In contrast, the numerosity of the other stimuli, including the target adaptor, the neutral adaptor, and the reference stimulus in the probe, was held constant at 40 dots throughout the experiment. We manipulated the contrast polarity configurations of the target adaptor. In the single-polarity condition, the contrast polarity of the dots in the target adaptor matched that of the other stimuli (i.e., the neutral adaptor and the test and the reference in the probe). In the mixed-polarity condition, the target adaptor consisted of an equal number of black and white dots.

We predicted that if numerosity perception is primarily adapted by dots sharing the same contrast polarity as the test stimulus, participants would overestimate numerosity in the mixed-polarity condition relative to the single-polarity condition. This overestimation would arise because only half of the dots in the mixed-polarity target adaptor (i.e., 20 dots) share the same contrast polarity as the test dot array.

Our results revealed a reliable difference in the points of subjective equality (PSEs) between single and mixed-polarity conditions (Figure 1B). Specifically, the PSE was significantly smaller in the mixed-polarity condition than in the single-polarity condition (39.3 versus 41.0 dots; t(15) = 4.53, p < 0.001, d = 1.13, paired t-test) (Figure 1C). This shift indicates that, despite identical adaptor numerosities, participants perceived the test stimulus as more numerous in the mixed-polarity condition than in the single-polarity condition, consistent with our prediction. We interpret this finding as evidence that numerosity adaptation is sensitive to contrast polarity. In particular, perceived numerosity appears to be most effectively adapted by the number of items that share the same low-level visual features (i.e., contrast polarity of dots).

### Experiment 2: Aftereffects persist under incongruent contrast polarity

Although our first experiment suggested the possibility that numerosity adaptation may be primarily driven by dots that share the same contrast polarity in the adaptor and test stimuli, it remained unclear whether numerosity aftereffects would be entirely eliminated when all dots differed in contrast polarity between the adaptor and test. To address this issue, Experiment 2 manipulated the congruency of contrast polarity between the adaptor and probe stimuli, such that contrast polarity was either the same (congruent) or different (incongruent) (Figure 2A). Adaptor numerosity was set to either large (80 dots), small (20 dots), or neutral (40 dots) and was manipulated across blocks. We predicted that if numerosity adaptation depended exclusively on matching contrast polarity, the negative aftereffects would be abolished when contrast polarity was incongruent between adaptor and probe stimuli.

**Figure 2.**
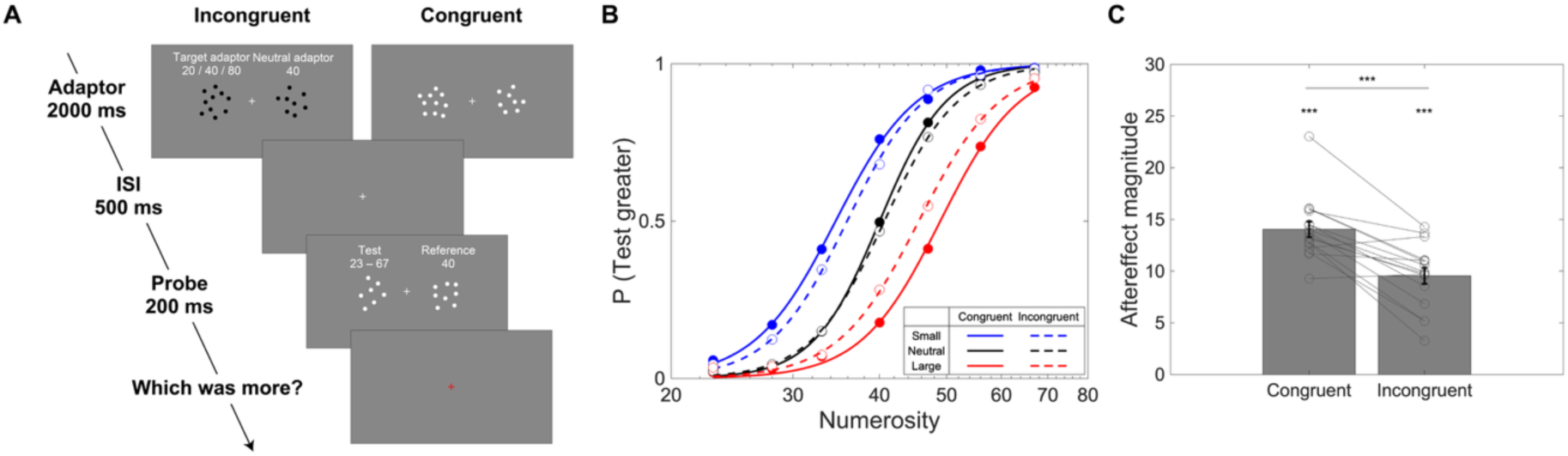
Stimulus sequence and results of Experiment 2. (A) As in Experiment 1, adaptor and probe stimuli were presented sequentially. The target adaptor contained either 20 (small), 40 (neutral), or 80 (large) dots, while the numerosity of the remaining stimuli (i.e., the neutral adaptor, test stimulus, and reference stimuli) was identical to that in Experiment 1. During the response period, participants indicated which of the two dot arrays in the probe stimulus contained more dots. For illustration purposes, fewer dots are shown than were used in the actual experiment. (B) Psychometric function fitted to the mean proportion of ‘test greater’ responses across participants for congruent (solid line) and incongruent (dashed line) contrast polarity conditions. Blue, red, and black lines represent the small (20 dots), large (80 dots), and neutral (40 dots) adaptation conditions, respectively. (C) Mean numerosity aftereffect magnitudes for congruent and incongruent contrast polarity conditions across participants. Gray circles represent individual participants’ data. Error bars denote standard error. *** p < 0.001.

A two-way repeated-measures ANOVA on PSEs revealed a significant interaction between adaptor numerosity and contrast polarity congruency (F(2, 30) = 17.40, p < 0.001, ηp² = 0.537), along with a significant main effect of adaptor numerosity (F(2, 30) = 187.31, p < 0.001, ηp² = 0.926) (Figure 2B). The main effect of congruency was not significant (F(1, 15) = 1.45, p = 0.248, ηp² = 0.088). These results indicate that perceived numerosity exhibited the expected repulsive aftereffects following adaptation to small and large numerosities, but that the magnitude of the aftereffect depended on whether contrast polarity was congruent or incongruent between adaptor and probe stimuli. This pattern was further confirmed by directly comparing aftereffect magnitudes, defined as the difference between PSEs following adaptation to large and small numerosities (Figure 2C). The aftereffect magnitude was significantly larger in the congruent condition (mean = 14.0 dots) than in the incongruent condition (mean = 9.5 dots; t(15) = 5.81, p < 0.001, d = 1.45, paired t-test). Critically, however, robust numerosity aftereffects remained evident even when contrast polarity was incongruent between adaptor and probe stimuli (congruent: t(15) = 18.81, p < 0.001, d = 4.70; incongruent: t(15) = 12.18, p < 0.001, d = 3.04; one-sample t-tests, p-values corrected for multiple comparisons). The persistence of robust aftereffects under incongruent conditions is inconsistent with an account in which numerosity adaptation depends exclusively on polarity-matched representations, indicating that feature-invariant adaptation contributes to the observed aftereffect.

We speculated that the asymmetry in aftereffect magnitude between the congruent and incongruent conditions may be attributable to low-level sensory adaptation to contrast polarity. Previous studies have proposed that the magnitude of contrast adaptation, defined as the difference between congruent and incongruent contrast-polarity conditions, is selective for contrast polarity and greater for darker than for lighter probe stimuli^31,32^. This predicts that if contrast adaptation at a low-level sensory-processing stage contributes to numerosity adaptation, the congruency effect in numerosity adaptation (i.e., aftereffect magnitude for congruent minus incongruent conditions) should be greater for black probes than for white probes, owing to stronger contrast adaptation when the probe has congruent black contrast polarity.

To explore this possibility, we split the data according to probe contrast polarity and examined the aftereffect magnitude in the congruent and incongruent conditions separately for each group (Figure 3). A two-way mixed ANOVA on aftereffect magnitudes revealed a significant main effect of congruency (F(1, 14) = 54.51, p < 0.001, ηp² = 0.796) and, crucially, a significant interaction between probe contrast polarity and congruency (F(1, 14) = 10.22, p = 0.006, ηp² = 0.422). Consistent with our prediction, the congruency effect was larger for black probes (6.43 dots) than for white probes (2.54 dots). These results support the idea that low-level contrast adaptation may contribute to numerosity aftereffects.

**Figure 3.**
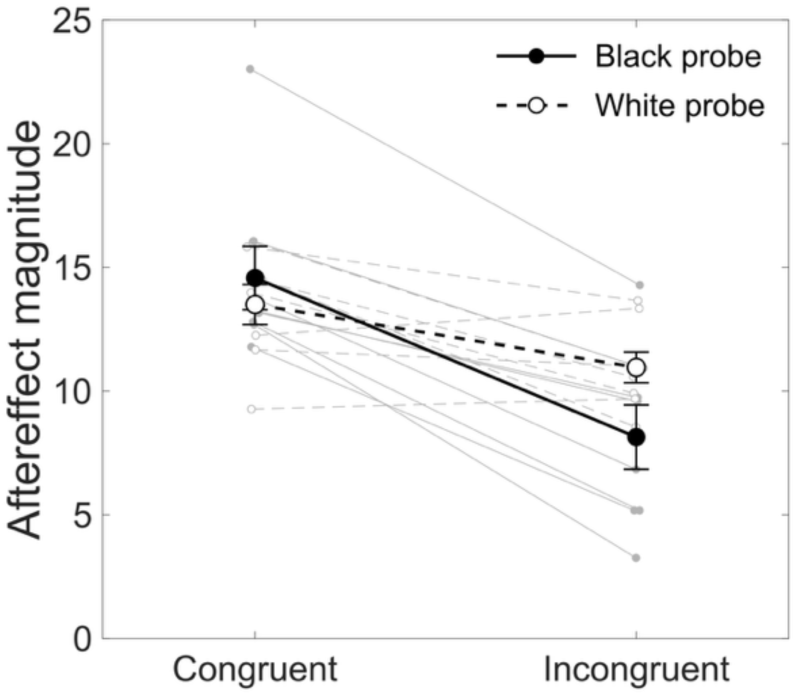
Aftereffect magnitude for black and white probes. Filled circles connected by a solid line represent the mean aftereffect magnitude in the congruent and incongruent conditions for the black-probe group (n = 8), whereas open circles connected by a dashed line represent the corresponding mean for the white-probe group (n = 8). Gray lines show individual participants’ data. Error bars denote standard error.

### Experiment 3: Afterimages do not account for residual aftereffects

These results demonstrate that although the magnitude of the adaptation effect is attenuated when contrast polarity is incongruent between the adaptor and test stimuli, numerosity aftereffects are not abolished. The persistence of aftereffects despite changes in contrast polarity suggests that numerosity aftereffects may arise from adaptation processes operating at multiple levels, including both high-level numerosity and low-level contrast polarity processing stages. However, an alternative explanation for the persistence of negative aftereffects in incongruent conditions involves retinal afterimages. Although the stimulus contrast in our experiments was relatively low, it is possible that participants experienced retinal afterimages following prolonged exposure to the adaptor stimuli. Because the perceived contrast polarity of a retinal afterimage is opposite to that of the original adaptor stimulus, participants in the incongruent condition may have been effectively adapted to a retinal afterimage that was congruent with the contrast polarity of the test stimulus. Such an effect could, in principle, contribute to the observed persistence of numerosity aftereffects in the incongruent condition.

To rule out this possibility, we modified the design of Experiment 2 to minimize the likelihood of afterimage perception in Experiment 3. Specifically, we further reduced the contrast of the dot stimuli (white: 32 cd/m², black: 12 cd/m², background: 22 cd/m²) and introduced a white-noise mask (500-ms duration) immediately following both the adaptor and probe stimuli (Figure 4A). Aside from these modifications, all other stimulus parameters were identical to those used in Experiment 2. Although we expected these changes to substantially reduce the perception of retinal afterimages, we additionally asked participants to rate the frequency and intensity of any perceived afterimages on a 7-point Likert scale after completing the experiment. This allowed us to directly assess the potential contribution of retinal afterimages to the observed numerosity aftereffects.

**Figure 4.**
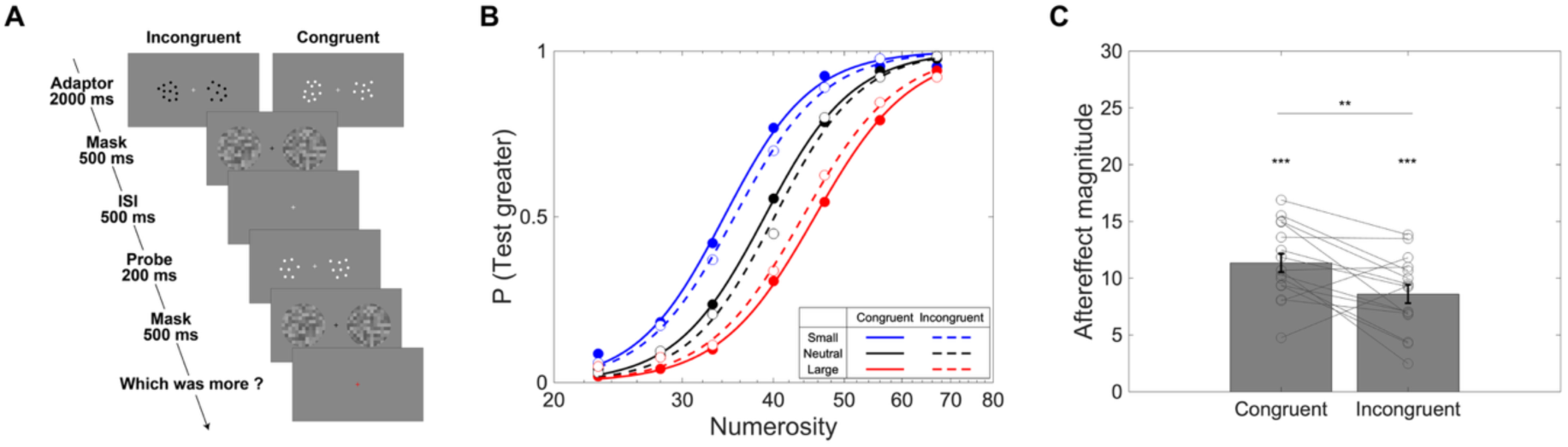
Stimulus sequence and results of Experiment 3. (A) Except for the insertion of a 500-ms mask stimulus immediately following the adaptor and probe stimuli, the stimulus sequence was identical to that of Experiment 2. (B) Psychometric functions fitted to the mean proportion of ‘test greater’ responses across participants. Solid and dashed lines indicate congruent and incongruent contrast polarity conditions, respectively. Blue, red, and black lines represent the small (20 dots), large (80 dots), and neutral (40 dots) adaptation conditions. (C) Mean numerosity aftereffect magnitudes for congruent and incongruent contrast polarity conditions across participants. Gray circles represent individual participants’ data. Error bars denote standard error. ** p < 0.01; *** p < 0.001.

A two-way repeated-measures ANOVA on PSEs again revealed a significant interaction between adaptor numerosity and contrast polarity congruency (F(2, 30) = 5.84, p = 0.007, ηp² = 0.280) along with a significant main effect of adaptor numerosity (F(2, 30) = 142.55, p < 0.001, ηp² = 0.905) (Figure 4B). Consistent with Experiment 2, the main effect of congruency was not significant (F(1, 15) = 0.01, p = 0.912, ηp² < 0.001). Direct comparison of aftereffect magnitude revealed that the aftereffect was significantly smaller in the incongruent condition (mean = 8.6 dots) than in the congruent condition (mean = 11.3 dots; t(15) = 3.02, p = 0.009, d = 0.75, paired t-test) (Figure 4C). Importantly, robust numerosity aftereffects were still observed in both conditions (congruent: t(15) = 13.98, p < 0.001, d = 3.50; incongruent: t(15) = 10.53, p < 0.001, d = 2.63; one-sample t-tests, p-values corrected for multiple comparisons). Crucially, Spearman’s rank correlations revealed no significant associations between aftereffect magnitude in the incongruent condition and subjective afterimage ratings for adaptor (intensity: ρ = 0.15, p = 1.000, Figure 5A; frequency: ρ = 0.05, p = 1.000, Figure 5B; p-values corrected for multiple comparisons) or for probe (intensity: ρ = -0.07, p = 1.000, Figure 5C; frequency: ρ = -0.12, p = 1.000, Figure 5D; p-values corrected for multiple comparisons). This finding argues against a substantial contribution of retinal afterimages to the observed aftereffect magnitudes. Together, these results replicate and extend the findings of Experiment 2 and indicate that retinal afterimages are unlikely to account for the persistence of numerosity aftereffects in incongruent contrast-polarity conditions.

**Figure 5.**
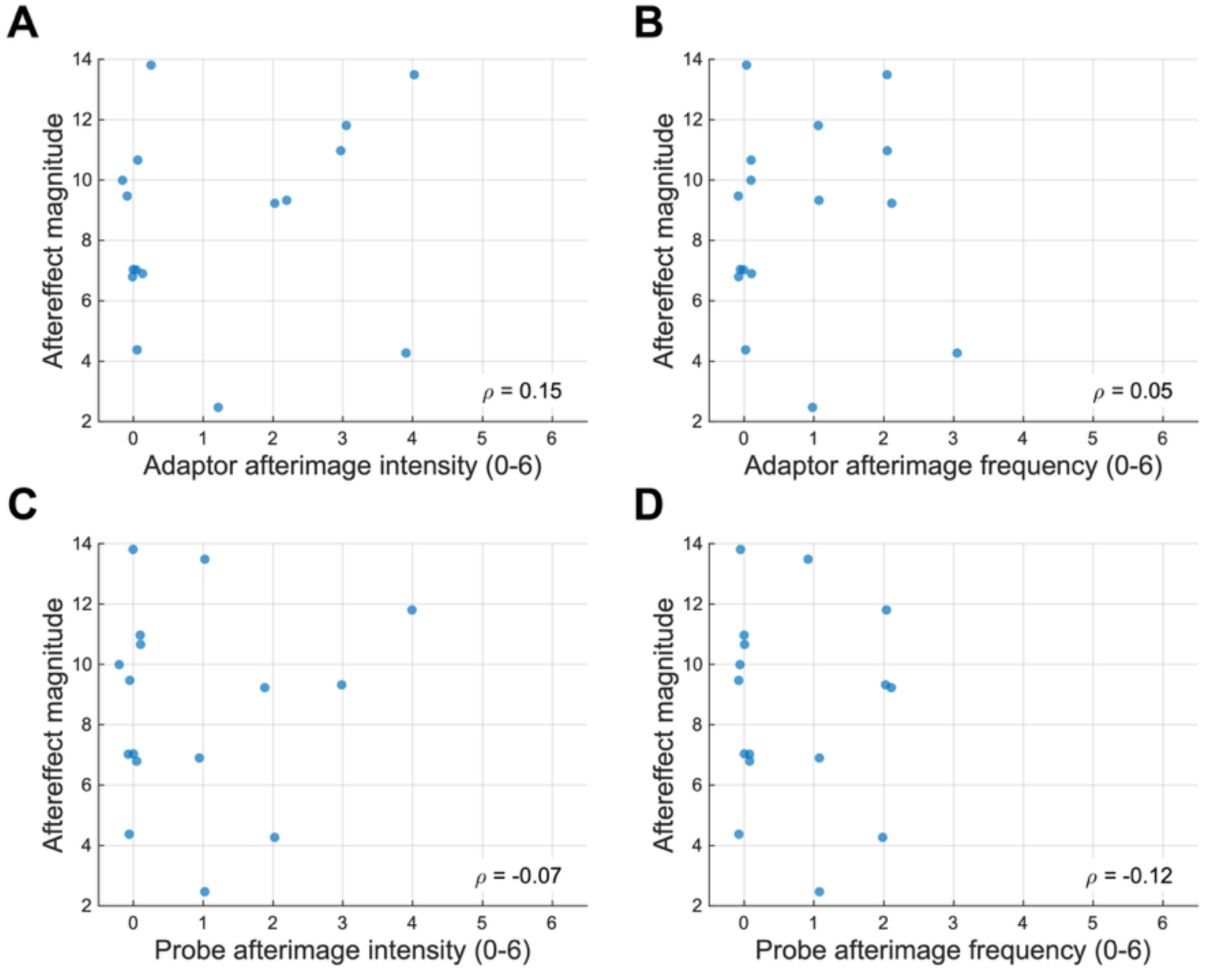
Correlations between numerosity aftereffect magnitude and Likert scale scores of afterimages in the incongruent condition. Correlation between aftereffect magnitude and rated intensity (A) and frequency (B) of afterimages following the adaptor stimulus. Correlation between aftereffect magnitude and rated intensity (C) and frequency (D) of afterimages following the probe stimulus. Each dot represents an individual participant. Spearman’s rank correlation coefficients (ρ) are shown in each panel.

### Experiment 4: Predictability of contrast polarity was not relevant to the residual aftereffects

Another possible account for the persistence of numerosity aftereffects in incongruent conditions concerns the predictability of the contrast polarity of the probe stimuli. In both Experiments 2 and 3, the contrast polarity of the adaptor stimuli varied across blocks, whereas the contrast polarity of the probe stimuli remained identical within each participant. This design allowed participants to accurately predict the contrast polarity of the upcoming probe stimuli. Previous research has shown that neural responses to numerosity in the superior parietal lobule depend on the attended contrast polarity^29^. Accordingly, we considered the possibility that the predictability of contrast polarity biased participants’ attention toward a predicted contrast polarity, thereby contributing to the apparent contrast-polarity invariance of numerosity adaptation in parietal cortex and, consequently, to the persistence of negative aftereffects in incongruent conditions.

To directly test whether predictability of contrast polarity contributes to the persistence of numerosity aftereffects, we conducted Experiment 4 in which the contrast polarity of the probe stimuli was rendered unpredictable by randomizing polarity on a trial-by-trial basis.

The results of Experiment 4 again confirmed robust negative numerosity aftereffects in both congruent and incongruent contrast polarity conditions (Figure 6A). A two-way repeated-measures ANOVA on PSEs revealed a significant main effect of adaptor numerosity (F(2, 30) = 180.80, p < 0.001, ηp² = 0.923) and a significant interaction between adaptor numerosity and contrast polarity congruency (F(2, 30) = 7.02, p = 0.003, ηp² = 0.319). Consistent with Experiments 2 and 3, the main effect of congruency was not significant (F(1, 15) = 1.40, p = 0.254, ηp² = 0.086).

**Figure 6.**
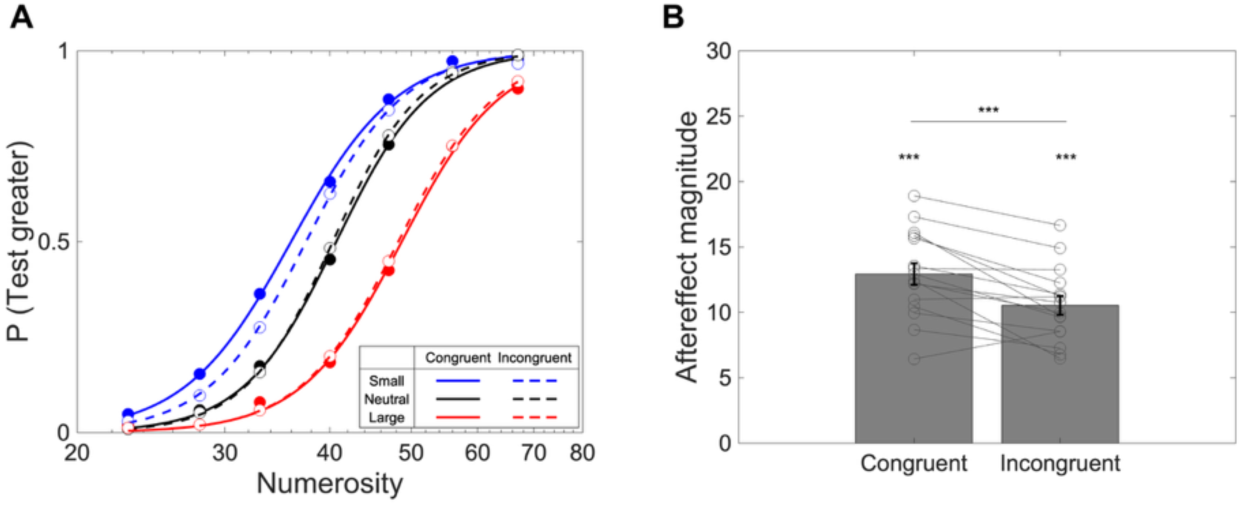
Results of Experiment 4. (A) Psychometric functions fitted to the mean proportion of ‘test greater’ responses across participants. Solid and dashed lines indicate congruent and incongruent contrast polarity conditions, respectively. Blue, red, and black lines represent the small (20 dots), large (80 dots), and neutral (40 dots) adaptation conditions. (B) Mean numerosity aftereffect magnitudes for congruent and incongruent contrast polarity conditions across participants. Gray circles represent individual participants’ data. Error bars denote standard error. *** p < 0.001.

We further replicated the pattern of aftereffect magnitudes observed in Experiments 2 and 3 (Figure 6B). The numerosity aftereffect was significantly larger in the congruent condition than in the incongruent condition (t(15) = 4.38, p < 0.001, d = 1.09, paired t-test). Critically, however, aftereffect magnitudes were significantly greater than zero in both conditions (congruent: t(15) = 15.74, p < 0.001, d = 3.94; incongruent: t(15) = 14.92, p < 0.001, d = 3.73; one-sample t-tests, p-values corrected for multiple comparisons).

To further examine whether the predictability of contrast polarity influenced the magnitude of numerosity aftereffects, we conducted a two-way mixed ANOVA with experiment (Experiment 2 vs. Experiment 4) as a between-subjects factor and congruency (congruent vs. incongruent) as a within-subjects factor. As expected, this analysis revealed a strong main effect of congruency (F(1, 30) = 52.86, p < 0.001, ηp² = 0.638), reflecting larger aftereffects in the congruent than in the incongruent condition. Although the interaction between Experiment and Congruency was statistically significant (F(1, 30) = 4.83, p = 0.036, ηp² = 0.139), post-hoc comparisons revealed no significant differences between Experiment 2 and Experiment 4 within either congruency condition (congruent: t(30) = 0.99, p = 0.660, d = 0.35; incongruent: t(30) = -0.93, p = 0.719, d = -0.33; two-sample t-tests, p-values corrected for multiple comparisons). These findings indicate that the numerosity aftereffects observed under incongruent conditions cannot be attributed to the predictability of probe polarity.

### Computational model with multi-level adaptation mechanisms reproduced behavioral results

Finally, we conducted computational simulations to examine whether the observed behavioral results could be accounted for by incorporating low-level sensory adaptation in addition to high-level numerosity-selective adaptation. The model comprised two mechanisms. First, a low-level, contrast-polarity-selective adaptation process that reduced the effective numerosity of the probe only when the adaptor and probe shared the same contrast polarity. Second, adaptation-driven gain reduction in numerosity-tuned neural populations, which affected the congruent and incongruent conditions equally (see Methods for details). We hypothesized that these two mechanisms would jointly produce larger aftereffects in the congruent condition than in the incongruent condition, while preserving robust aftereffects in the incongruent condition.

The simulations showed that incorporating low-level contrast-polarity-selective adaptation, in addition to high-level numerosity adaptation, qualitatively reproduced the principal behavioral observations. When no contrast-polarity-selective contrast adaptation was assumed in either the congruent or incongruent condition (i.e., when both the congruent and incongruent ratios were set to 0; see Methods for details), no systematic difference in the shifts of the psychometric functions was observed between the two conditions (Figure 7A). In contrast, when contrast-polarity-selective contrast adaptation was assumed to reduce the effective numerosity in the probe stimulus (i.e., when the congruent ratio was increased to 0.05), the magnitude of the shifts differed between conditions (Figure 7B). In the congruent condition, early-stage adaptation reduced the effective numerosity of both the test and reference stimuli, thereby increasing the apparent magnitude of the aftereffect. In the incongruent condition, by contrast, this early-stage adaptation effect was absent, and the observed shifts reflected the influence of numerosity-selective adaptation alone. These results suggest that the combined effects of low-level contrast polarity-selective contrast adaptation and high-level numerosity-tuned adaptation can account for our behavioral observation that aftereffect magnitudes were significantly reduced, yet remained robust, when low-level sensory features were mismatched.

**Figure 7.**
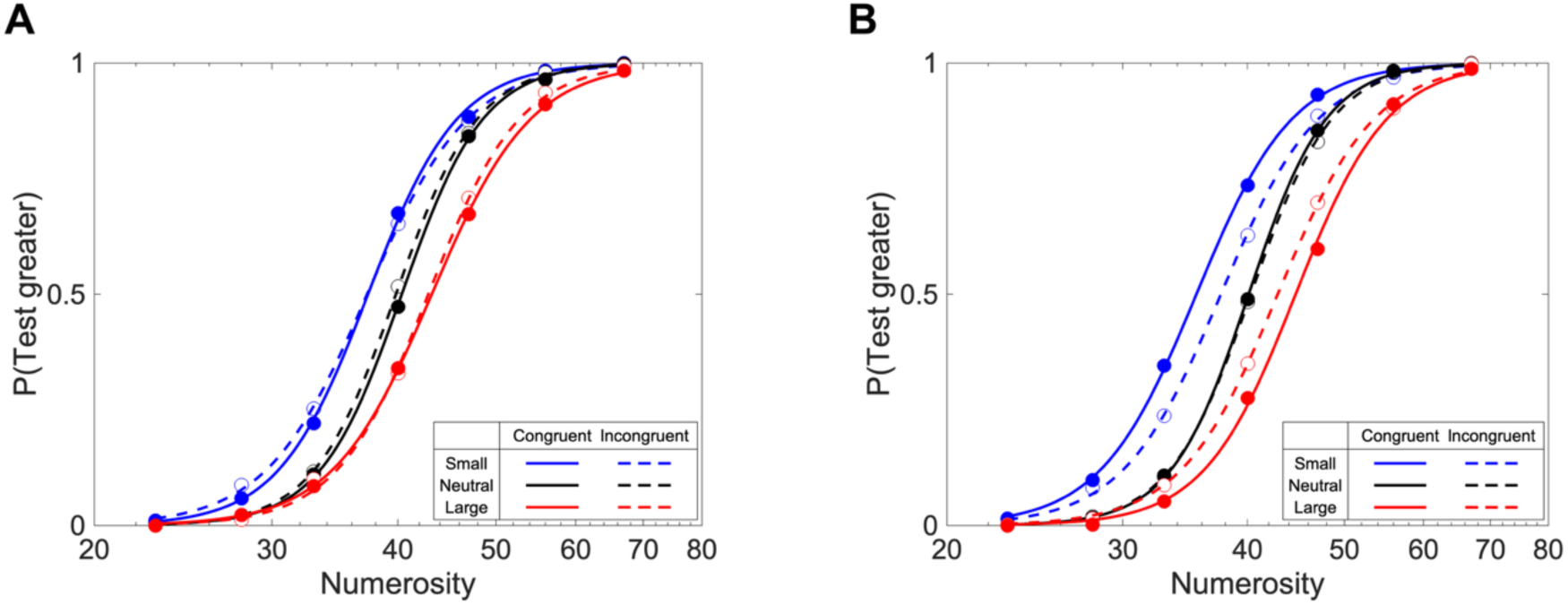
Results of computational simulations. (A) Simulation results with both the congruent and incongruent ratio set to 0. (B) Simulation results with a congruent ratio of 5% and an incongruent ratio of 0. Solid lines indicate the congruent contrast polarity condition, and dashed lines indicate the incongruent condition. Blue, red, and black lines represent the small, large, and neutral adaptor conditions, respectively.

## Discussion

We investigated how contrast-polarity congruency between adaptor and test stimuli modulates numerosity adaptation, with the aim of distinguishing two possible accounts of feature sensitivity: a direct effect, in which high-level numerosity representations are themselves selective for low-level visual features, and an indirect effect, in which earlier sensory adaptation modulates the input to otherwise feature-invariant numerosity representations. We predicted that, if feature sensitivity reflects a direct effect, numerosity aftereffects should be observed only when contrast polarity is congruent between adaptor and test stimuli. In contrast, if feature sensitivity arises from an indirect effect, the aftereffect should be reduced but should persist under incongruent conditions. Our results support the indirect account. Across a series of psychophysical experiments, we consistently showed that switching contrast polarity significantly reduced the magnitude of the aftereffect, yet robust negative aftereffects persisted even when retinal afterimages and polarity predictability were controlled. We further showed that a computational model combining feature-invariant numerosity adaptation with contrast-polarity-selective sensory adaptation qualitatively reproduced this behavioral pattern. Together, these results suggest that sensitivity to contrast-polarity congruency in numerosity adaptation arises through an indirect effect.

The larger numerosity aftereffects observed in the congruent condition than in the incongruent condition parallel previously reported asymmetries in contrast adaptation. In a previous psychophysical study, Sato and colleagues showed that the perceived contrast of sparsely distributed elements, similar to the stimuli used in the present study, was significantly reduced when the contrast polarity of the test stimulus matched that of the adaptor, compared with when contrast polarity was unmatched^31^. Importantly, they also reported that this effect interacted with contrast polarity, such that the congruency effect was larger for test stimuli with negative contrast polarity than for those with positive contrast polarity. A similar asymmetry was present in the numerosity aftereffects observed in our data: numerosity aftereffect magnitudes were generally larger for probes with negative contrast polarity than for those with positive contrast polarity. These results suggest that asymmetries in early sensory adaptation contribute to the larger numerosity aftereffects observed under congruent contrast-polarity conditions.

Based on these phenomenological similarities, we propose that the reduced but significant residual aftereffect in the incongruent condition, relative to the congruent condition, reflects adaptation at two functionally distinct processing stages: a numerosity-processing stage and a sensory-processing stage. The numerosity-processing stage comprises feature-invariant, numerosity-tuned representations and is therefore susceptible to adaptation to numerosity itself. In contrast, the sensory-processing stage provides the input to the higher-level numerosity processing stage, such that adaptation at either stage can influence the magnitude of the observed numerosity aftereffect.

How might asymmetry in early sensory adaptation influence numerosity aftereffects at a later processing stage? One possible physiological implementation of the proposed sensory-processing stage is that asymmetric contrast adaptation alters aggregate Fourier power, which has been proposed as a potential source of numerosity information^33,34^ that is subsequently extracted by tuned, high-level numerosity representations^27,28^. When adaptor and test dot arrays share the same contrast polarity, adaptation in low-level spatial-frequency channels would reduce the effective sensory input driven by the test dot arrays, leading to a decrease in aggregate Fourier power. Because the reduction in aggregate Fourier power would be expected to scale with adaptor numerosity, adaptation to a large numerosity should produce a greater reduction than adaptation to a small numerosity. Since the neutral adaptor had an intermediate numerosity and was paired with the reference stimulus, it should produce an intermediate reduction in aggregate Fourier power of the reference stimulus. These modulations of aggregate Fourier power could bias inputs to the numerosity-processing stage and consequently influence numerosity judgements, in the same direction as the numerosity aftereffect: compared with the reference, numerosity would tend to be judged as smaller following adaptation to a large numerosity and as larger following adaptation to a small numerosity. In contrast, when adaptor and probe stimuli were incongruent, polarity-selective sensory adaptation would be expected to produce little or no reduction in aggregate Fourier power.

Importantly, in our proposed framework, asymmetries in aggregate Fourier power would add to feature-invariant numerosity adaptation at the numerosity-processing stage. This provides a potential explanation for the persistence of numerosity aftereffects when contrast polarities were unmatched: although reversing contrast polarity between adaptor and probe stimuli abolishes, or at least substantially reduces, the contribution of contrast adaptation to the numerosity readout, the aftereffect persists because of genuine, feature-invariant numerosity adaptation at the numerosity-processing stage. Consistent with this interpretation, our phenomenological computational model, which incorporated low-level sensory adaptation alongside feature-invariant numerosity adaptation, qualitatively reproduced the observed behavioral pattern.

Although our two-stage model acknowledges the contribution of low-level sensory adaptation to numerosity aftereffects, it challenges the notion that numerosity adaptation can be attributed entirely to low-level sensory adaptation, as proposed by the old news hypothesis^35,36^. Across experiments, we showed that aftereffects persisted when adaptor and probe stimuli had incongruent contrast polarity, a condition where polarity-sensitive contrast adaptation would be expected to contribute little or no effect. This finding indicates that low-level sensory adaptation alone is insufficient to fully account for the numerosity aftereffect. Within our proposed framework, the aftereffect observed under incongruent conditions reflects feature-invariant numerosity adaptation, whereas the larger aftereffect observed under congruent conditions reflects the combined influence of feature-invariant numerosity adaptation and low-level sensory adaptation. Accordingly, feature-invariant numerosity adaptation appears to account for most of the observed aftereffect, with low-level sensory adaptation providing an additional, but comparatively modest, contribution. Together, these findings suggest that numerosity aftereffects are driven primarily by adaptation within feature-invariant numerosity representations, while also being shaped by earlier sensory adaptation.

A more recent study by Grasso and colleagues^37^ has also argued against the old-news account. By manipulating whether overlapping dot colors were matched or unmatched between adaptor and test stimuli, they showed that adaptation of tuned numerosity mechanisms alone better explains numerosity aftereffects than does the old-news account. However, they did not explicitly test the two-stage account proposed here. Specifically, our framework assumes that congruency in low-level sensory features modulates the overall sensory input to the numerosity processing stage, rather than the responses of individual sensory channels tuned to precise spatial locations, as proposed by Grasso and colleagues. From this perspective, their experimental findings are compatible with our two-stage account. Consistent with this interpretation, our phenomenological model incorporated an ensemble-level modulatory stage and showed that it can account for the behavioral pattern we observed without invoking position-level dot tracking.

Our two-stage account may also help explain an earlier finding on numerosity-adaptation that is difficult to reconcile with a purely tuned mechanism^13^. Burr and Ross^38^ reported that adaptation to a large number of dots reduced perceived numerosity. Notably, this aftereffect remained substantial even when the adaptor numerosity was far greater than the test numerosities. For example, adaptation to 400 dots reduced perceived numerosity by approximately half even for test displays containing only ∼12 dots, more than an order of magnitude fewer than the adaptor. Burr and Ross attributed this long-range aftereffect to broadly tuned numerosity-selective neurons. However, although numerosity tuning is broad, it is approximately scale-invariant on a logarithmic axis and corresponds to a neuronal Weber fraction of only ∼0.2–0.3^39,40^. An adaptor that differs so greatly from the test numerosity should therefore fall well outside the range of channels activated by the test. Consequently, genuine tuned adaptation would be expected to make only a limited contribution at such large adaptor– test separations. Within our framework, the aftereffect that persists across these large separations is instead attributed to low-level sensory adaptation. Its magnitude increases with adaptor numerosity because larger adaptors produce stronger contrast adaptation and a greater reduction in sensory energy. Importantly, this mechanism does not depend on overlap between numerosity-tuned channels. This account leads to a direct prediction. If the tuned component is isolated, for example, by using incongruent contrast polarity to minimize low-level adaptation, the aftereffect should be restricted to a narrower range of adaptor–test numerosity differences.

Our findings are largely consistent with the recent report by Caponi and colleagues^41^, which showed that aftereffect magnitudes depend on image dissimilarity across various sensory features, such as color, luminance, shape, and motion. They attributed differences in aftereffect magnitude between matched and unmatched conditions to high-level interactions, assuming the existence of numerosity representations for categorized features. However, it remains unclear how this high-level account can explain both the persistence of the aftereffect under incongruent contrast-polarity conditions and the asymmetric contrast adaptation between contrast polarities reported here. By focusing on contrast-polarity congruency, we instead suggest that feature sensitivity may arise as a consequence of low-level adaptation that indirectly modulates high-level numerosity processing, possibly through sensory energy information. Although further study is required, our findings suggest that feature sensitivity in numerosity adaptation is unlikely to arise solely from either low-level sensory adaptation or feature-specific numerosity representations. Instead, they support the view that low-level sensory adaptation interacts with feature-invariant numerosity processing to shape the observed aftereffects. This interpretation accounts for both the persistence of aftereffects under incongruent conditions and the larger aftereffects observed under congruent conditions, without requiring separate numerosity representations for different sensory features.

In conclusion, the present findings show that numerosity adaptation reflects adaptation across multiple stages of visual processing, rather than being explained solely by either low- or high-level mechanisms. Based on the results of psychophysical experiments and modeling, we propose a two-stage framework in which adaptation operates through independent mechanisms: contrast-polarity-selective adaptation at a low-level sensory-processing stage, and feature-invariant numerosity adaptation at a numerosity-processing stage. The effect of contrast adaptation at the sensory-processing stage might be conveyed as sensory energy to the numerosity-processing stage, where adaptation to numerosity itself occurs. This hierarchical structure allows numerosity-driven aftereffects to be modulated by low-level contrast adaptation. Together, our results provide important insight into how the visual system reconciles the vulnerability of numerosity aftereffects to feature changes with their persistence despite such changes.

## Limitations of the study

Several limitations should be noted. First, we examined only contrast polarity as the feature dimension; whether our two-stage model generalizes to other features, such as color, shape, or motion, remains to be tested. Second, our experiments used a fixed set of numerosities within the numerosity range^18,42^, and the generality of the model across wider numerosity ranges, including the subitizing range and the texture range, remains to be examined. Finally, our experiments did not dissociate spatiotopic and retinotopic reference frames, leaving open the question of whether low-level sensory adaptation and high-level numerosity adaptation operate in distinct coordinate systems.

## STAR Methods

### Key resources table

| REAGENT or RESOURCE | SOURCE | IDENTIFIER |
| --- | --- | --- |
| **Deposited data** |  |  |
| Source data for all figures | This paper | Zenodo:<br>https://doi.org/10.5281/zenodo.21585587 |
| **Software and algorithms** |  |  |
| MATLAB R2021b | MathWorks | https://it.mathworks.com/products/matlab.html |
| Psychtoolbox-3 (3.0.19) | Brainard, 1997; Pelli, 1997; Kleiner et al., 2007 | http://psychtoolbox.org/ |
| Palamedes 1.11.12 | Prins and Kingdom, 2018 | http://www.palamedestoolbox.org/ |
| JASP 0.19.3 | The JASP Team | http://jasp-stats.org/ |
| Simulation code | This paper | Zenodo:<br>https://doi.org/10.5281/zenodo.21585587 |

### Resource availability

#### Lead contact

Further information and requests for resources and reagents should be directed to and will be fulfilled by the lead contact, Masamichi J. Hayashi.

#### Materials availability

This study did not generate new unique reagents.

#### Data and code availability

- The source data underlying all figures in this paper have been deposited at Zenodo and are publicly available. The DOI is listed in the key resources table. All other data reported in this paper are available from the lead contact upon request.
- The simulation code used to generate Figure 7 has been deposited at Zenodo and is publicly available. The DOI is listed in the key resources table. Any other original code is available from the lead contact upon request.
- Any additional information required to reanalyze the data reported in this paper is available from the lead contact upon request.

### Experimental model and study participant details

#### Participants

Sixty-four healthy adult volunteers participated in the experiments (Experiment 1: 12 males and 4 females, mean age = 23.6 ± 1.2 years; Experiment 2: 10 males and 6 females, mean age = 22.0 ± 1.6 years; Experiment 3: 10 males and 6 females, mean age = 22.9 ± 1.5 years; Experiment 4: 11 males and 5 females, mean age = 21.9 ± 1.3 years). All participants were right-handed, had normal or corrected-to-normal vision, and reported no history of neuropsychiatric disorders. The study was conducted in accordance with the Declaration of Helsinki. All participants provided written informed consent prior to participation. The experimental protocol was approved by the institutional ethics committee of the National Institute of Information and Communications Technology.

### Method details

#### Apparatus

Stimulus generation, experimental control, and response collection were implemented in MATLAB R2021b (MathWorks, Natick, MA, USA) using Psychophysics Toolbox Version 3.0.19^43–45^, running on Ubuntu 22.04 LTS. Stimuli were presented on a gamma-corrected VIEWPixx/3D monitor (22.5-inch, resolution = 1920 × 1200 pixels, refresh rate = 100 Hz, VPixx Technologies, Saint-Bruno, QC, Canada). Stimulus timing was validated using an oscilloscope. Participants viewed the display from a distance of 57 cm, with head position stabilized using a chin rest. Responses were collected using a standard keyboard.

#### Experimental design and procedure

The stimulus sequence and configurations were essentially identical across all experiments. On each trial, an adaptor stimulus (2,000 ms) and a probe stimulus (200 ms) were presented sequentially on a gray background, separated by a 500-ms interstimulus interval (ISI). The adaptor stimulus comprised two dot arrays: a target adaptor and a neutral adaptor, presented in the left and right visual fields (one on each side). The probe stimulus consisted of a test dot array presented at the same location as the preceding target adaptor and a reference dot array presented at the same location as the preceding neutral adaptor. At probe offset, the central fixation cross turned red to indicate the response period. Participants were instructed to compare the two dot arrays in the probe (test vs. reference) and indicate which array contained more dots (two-alternative forced-choice task). The next trial began immediately after a response was recorded.

In both adaptor and probe stimuli, dot arrays were displayed within a virtual circular region (radius = 6°) centered 8° to the left and right of a central fixation cross. Individual dots subtended 0.2° in diameter, and a minimum center-to-center distance of 1° was maintained between dots. Across all experiments, adaptor and probe numerosities were chosen to fall within the numerosity range, because numerosities in other ranges, such as the subitizing or texture-density ranges, are thought to rely on distinct perceptual mechanisms^46–49^.

##### Experiment 1

To examine the influence of mixing dots with opposite contrast polarities, Experiment 1 manipulated the contrast polarity of the adaptor stimulus against a gray background (22 cd/m²) under two conditions: a single-polarity condition and a mixed-polarity condition (Figure 1A). In the single-polarity condition, all dot arrays in both the adaptor and probe stimuli had the same contrast polarity, either white (42 cd/m²) or black (2 cd/m²). Each participant completed both the white and black conditions (i.e., contrast polarity was manipulated within participants). In the mixed-polarity condition, the target adaptor contained equal numbers of white and black dots, whereas the other dot arrays (i.e., the neutral adaptor in the adaptor stimulus and the test and reference dot arrays in the probe stimulus) had the same contrast polarity as in the single-polarity condition.

In both conditions, the numerosities of the target and neutral adaptors in the adaptor stimulus and of the reference dot array in the probe stimulus were fixed at 40 dots. By contrast, the numerosity of the test dot array varied across seven levels (23, 28, 33, 40, 47, 56, or 67 dots), each presented in 28 trials in each of the four condition-by-polarity combinations, in random order. The order of the condition-by-polarity combinations and the visual field in which the target and neutral adaptors appeared were counterbalanced across participants.

The experiment comprised eight blocks of 98 trials each. To minimize fatigue, participants took a one-minute break between blocks and a longer five-minute break after completing the first half of the experiment (i.e., after the fourth block).

#### Experiment 2

Experiment 2 further examined the effect of contrast-polarity congruency between adaptor and probe stimuli. Whereas Experiment 1 used dot arrays with mixed contrast polarities (i.e., equal numbers of white and black dots), Experiment 2 defined contrast polarity at the dot-array level (i.e., all dots within a given array were either white or black). We manipulated contrast polarity congruency such that the contrast polarity of the adaptor and probe stimuli was either matched (congruent) or mismatched (incongruent) (Figure 2A).

In each block, one of three adaptor numerosities, small (20 dots), neutral (40 dots), or large (80 dots), was used for the target adaptor and presented together with a neutral adaptor (40 dots). The reference numerosity was fixed at 40 dots, whereas the test numerosity varied across seven levels (23, 28, 33, 40, 47, 56, or 67 dots) in random order across trials. Thus, the experiment followed a 2 (congruent vs. incongruent) × 3 (small vs. neutral vs. large target-adaptor numerosity) within-subjects design, comprising 1,176 trials delivered in 12 blocks of 98 trials.

The experiment comprised 12 blocks of 98 trials each, organized into two halves separated by a five-minute break. Each half consisted of two baseline blocks (neutral target adaptor, 40 dots) followed by four adaptation blocks in which small-numerosity (20 dots) and large-numerosity (80 dots) target adaptors alternated (e.g., small, large, small, large). Block-to-block breaks were one minute.

The contrast polarity of the probe stimulus (white or black) was fixed for each participant and counterbalanced across participants. The contrast polarity of the adaptor stimulus (white or black) was varied across blocks within each participant; because the probe contrast polarity was fixed, this produced both congruent and incongruent conditions for each participant. Four binary factors were counterbalanced across participants: (1) the contrast polarity of the probe stimulus (white or black); (2) the visual field of the target adaptor (left or right); (3) the alternation order of adaptor numerosity across halves; and (4) the alternation order of adaptor polarity across blocks. These factors yielded 16 unique combinations, and each of the 16 participants was randomly assigned to one. The luminance values for white dots, black dots, and the gray background were identical to those in Experiment 1.

#### Experiment 3

Experiment 3 was designed to rule out the possibility that the persistence of numerosity aftereffects under contrast-polarity incongruence observed in Experiment 2 could be explained by retinal afterimages induced by high-contrast stimulus presentations.

The experimental design and stimulus configuration were identical to those of Experiment 2, except for the following modifications. First, stimulus contrast was reduced by adjusting luminance values (white dots, 32 cd/m²; black dots, 12 cd/m²; gray background, 22 cd/m²). Second, a 500-ms white-noise mask was inserted immediately after the offsets of both the adaptor and probe stimuli. Finally, after completing the experiment, participants rated the frequency and intensity of perceived afterimages using a 7-point Likert scale.

#### Experiment 4

Experiment 4 tested whether the persistence of numerosity aftereffects in incongruent conditions could be attributed to the predictability of probe contrast polarity. The experimental design and stimulus configuration were identical to those of Experiment 2, except that probe contrast polarity was randomized on a trial-by-trial basis, rendering probe contrast polarity unpredictable.

### Simulation

To examine whether our behavioral findings could be explained by a combination of low-level sensory adaptation and feature invariant numerosity adaptation, we constructed a computational model and assessed whether it could qualitatively reproduce the behavioral results observed in Experiments 2–4. The specific constraints used to construct the model are described below.

#### Neuronal population and tuning

We simulated a population of 500 model neurons with logarithmically spaced preferred numerosities between 1 and 250. Each unit’s response profile was modeled as a log-Gaussian tuning curve with σ = 0.4 in natural-log units, scaled to a maximum response of 0.18, motivated by evidence for numerosity-selective neurons^28,39,50,51^.

#### Feature invariant numerosity adaptation

Adaptation was implemented as a multiplicative gain change centered on the adaptor numerosity (20, 40, or 80), consistent with neuroimaging evidence that adaptation modulates numerosity-selective responses^27,29,52,53^ and following channel-based models of sensory adaptation^54,55^. The gain profile was log-Gaussian, matching the tuning bandwidth (σ = 0.4), with a maximum gain reduction of 50%. For the test stimulus, the gain profile reduced the responses of units with preferred numerosities close to the adaptor. For the reference stimulus (fixed at 40 dots), an analogous gain reduction profile was always applied, centered on 40 dots.

#### Low-level contrast-polarity-selective adaptation

We introduced an additional low-level sensory adaptation process that selectively affected the congruent condition, motivated by polarity-selective contrast adaptation. In the congruent condition, the numerosity used to generate the population responses for both test and reference dot arrays was reduced by an amount proportional to adaptor numerosity (congruency ratio, set to 5% of adapting numerosity). In the incongruent condition, no such reduction was applied (incongruent ratio, set to zero), such that responses were influenced only by adaptation-driven gain changes in the numerosity-tuned population described above. This implements the old news hypothesis^35^: early-stage underestimation processes (e.g., contrast-polarity-selective adaptation^32^) jointly affect the test and reference when the stimuli share the same contrast polarity (congruent), but not when they differ (incongruent).

#### Trial generation and decoding

On each trial, spike counts were sampled independently from Poisson distributions with means given by the adapted population responses. The resulting population activity was decoded using fixed, unadapted response templates. For each candidate numerosity *x*, the decoder computed the score:

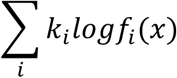

where *k_i_* is the sampled spike count of the *i*-th model neuron and *f_i_*(*x*) is the unadapted response of that neuron to candidate numerosity *x* . The candidate numerosity with the maximum score was selected as the decoded estimate.

## Choice rule and psychometric analysis

A binary choice was generated by comparing the decoded estimates of the test and reference numerosities. Each condition (congruent vs. incongruent) and adaptor numerosity (20, 40, or 80) was simulated for 16 observers, with 30 trials per test numerosity level. For each condition, responses were averaged across the 16 simulated observers, and a logistic function of log2 (test numerosity) was fit to this group-averaged psychometric curve, yielding a single PSE and slope per condition.

## Quantification and statistical analysis

For each participant, trials with response times exceeding the participant’s mean ± 3 SD were excluded from analysis (mean ± SD across participants: Experiment 1, 1.86 ± 0.63%; Experiment 2, 1.60 ± 0.59%; Experiment 3, 1.42 ± 0.68%; Experiment 4, 1.72 ± 0.59%). For each condition, the proportion of trials in which the test dot array was judged more numerous than the reference dot array was fit as a function of test numerosity using a logistic psychometric function in log2 space, implemented in the Palamedes toolbox^56^. From the fitted psychometric function, we extracted each participant’s PSE, defined as the test numerosity corresponding to a 50% probability of judging the test as more numerous than the reference (i.e., the perceived numerosity matches the reference numerosity). Effect sizes are reported as Cohen’s d for t-tests and partial eta-squared (ηp²) for ANOVAs.

In Experiment 1, we tested whether mixing dots with opposite contrast polarities induced a bias in perceived numerosity by comparing PSEs between the single- and mixed-polarity conditions using a two-tailed paired t-test (α = 0.05).

In Experiments 2–4, we tested for repulsive aftereffects using a two-way repeated-measures ANOVA on PSEs with within-subjects factors of contrast-polarity congruency (congruent vs. incongruent) and adaptor numerosity (small, neutral, and large). Aftereffect magnitude was computed as the PSE in the large-adaptor condition minus the PSE in the small-adaptor condition. A two-tailed paired t-test (α = 0.05) was used to assess whether aftereffect magnitude differed between the congruent and incongruent conditions. In addition, one-sample t-tests (two-tailed, α = 0.05) were used to test whether aftereffect magnitudes were significant within each congruency condition. For the two one-sample tests within each experiment, p-values were corrected for multiple comparisons using the Bonferroni method.

In Experiment 2, we further tested whether the effect of congruency on aftereffect magnitude depended on probe contrast polarity, using a two-way mixed ANOVA on aftereffect magnitudes with probe contrast polarity (black vs. white) as a between-subjects factor and congruency (congruent vs. incongruent) as a within-subjects factor (α = 0.05).

In Experiment 3, we additionally examined whether individual differences in aftereffect magnitude were associated with subjective reports of retinal afterimages. We computed Spearman’s rank correlations between aftereffect magnitude in the incongruent condition and afterimage ratings for intensity and frequency. Statistical significance was evaluated at α = 0.05 with Bonferroni correction for the four correlations.

We also tested whether the predictability of contrast polarity influenced the magnitude of numerosity aftereffects by directly comparing aftereffect magnitudes between Experiment 2 and Experiment 4. We conducted a two-way mixed ANOVA with experiment (Experiment 2 vs. Experiment 4) as a between-subjects factor and congruency (congruent vs. incongruent) as a within-subjects factor (α = 0.05). Post-hoc pairwise comparisons were conducted with Bonferroni correction.

## Acknowledgments

We thank Hiroshi Ban for his helpful advice on the experimental design. This work was supported by the Japan Society for the Promotion of Science (Grants-in-Aid for Scientific Research JP21H00315, JP23K22381, and JP23K17649 to MJH) and the Japan Science and Technology Agency (FOREST JPMJFR232X to MJH).

## Author contributions

Conceptualization, T.J.T. and M.J.H.; Methodology, T.J.T. and M.J.H.; Investigation, T.J.T. and N.W.R.; Formal analysis, T.J.T.; Writing – original draft, T.J.T.; Writing – review & editing, T.J.T., N.W.R., and M.J.H.; Supervision, M.J.H.

## Declaration of generative AI and AI-assisted technologies in the manuscript preparation process

During the preparation of this work, the authors used ChatGPT for improving the readability and language of the manuscript. The authors reviewed and edited the output as needed and take full responsibility for the content of the published article.

